# Analyses of Neanderthal introgression suggest that Levantine and southern Arabian populations have a shared population history

**DOI:** 10.1101/438390

**Authors:** Deven N. Vyas, Connie J. Mulligan

**Author notes:** Present Address: Department of Ecology and Evolution, Stony Brook University, Stony Brook, New York (Deven N. Vyas). Corresponding author: Deven N. Vyas, Department of Ecology & Evolution, Stony Brook University, 650 Life Sciences Building, Stony Brook, NY 11794-5245.

## Abstract

**Objectives:** Modern humans are thought to have interbred with Neanderthals in the Near East soon after modern humans dispersed out of Africa. This introgression event likely took place in either the Levant or southern Arabian depending on which dispersal route out of Africa was followed. In this study, we compare Neanderthal introgression in contemporary Levantine and southern Arabian populations to investigate Neanderthal introgression and to study Near Eastern population history.

**Materials and Methods:** We analyzed genotyping data on >400,000 autosomal SNPs from seven Levantine and five southern Arabian populations and compared those data to populations from around the world including Neanderthal and Denisovan genomes. We used f_4_ and D statistics to estimate and compare levels of Neanderthal introgression between Levantine, southern Arabian, and comparative global populations. We also identified 1,581 putative Neanderthal-introgressed SNPs within our dataset and analyzed their allele frequencies as a means to compare introgression patterns in Levantine and southern Arabian genomes.

**Results:** We find that Levantine and southern Arabian populations have similar levels of Neanderthal introgression to each other but lower levels than other non-Africans. Furthermore, we find that introgressed SNPs have very similar allele frequencies in the Levant and southern Arabia, which indicates that Neanderthal introgression is similarly distributed in Levantine and southern Arabian genomes.

**Discussion:** We infer that the ancestors of contemporary Levantine and southern Arabian populations received Neanderthal introgression prior to separating from each other and that there has been extensive gene flow between these populations.

---

Anatomically modern humans first evolved in Africa around 200 thousand years ago (kya), and later dispersed out of Africa (OOA) to settle the rest of the world (e.g., Cann, Stoneking, & Wilson, 1987; McDougall, Brown, & Fleagle, 2005; Stringer & Andrews, 1988; Stringer, 2002; White et al., 2003). Analyses of Neanderthal genomes suggest that all living people of non-African ancestry have low levels of Neanderthal introgression in their genomes, generally ranging around 1.5-2.6%. These low levels of Neanderthal introgression are thought to reflect contact between Neanderthals and early modern humans following the OOA dispersal (Green et al., 2010; Prüfer et al., 2017; Prüfer et al., 2014). Genomic studies have consistently dated the main OOA dispersal (i.e., the dispersal from which the overwhelming majority of non-African genetic diversity derives) to ~50-70 kya (e.g., Malaspinas et al., 2016; Mallick et al., 2016; Pagani et al., 2016). Correspondingly, introgression has been dated to ~50-60 kya (Fu et al., 2014; Sankararaman, Patterson, Li, Pääbo, & Reich, 2012; Seguin-Orlando et al., 2014), which is remarkably consistent with the dating of the main OOA dispersal. Given the consistency of dates for the main OOA dispersal and Neanderthal introgression, as well as the fact that introgression is present in all non-Africans, it has been proposed that the introgression event occurred very soon after the OOA dispersal and most likely somewhere in the Near East (e.g., Alves, Šrámková Hanulová, Foll, & Excoffier, 2012; Currat & Excoffier, 2011; Green et al., 2010; Lazaridis et al., 2016; Prüfer et al., 2014; Wall et al., 2013).

Despite Neanderthal introgression being present in all non-African populations, populations from different parts of the world have different levels of introgression, with the highest levels in East Asians (e.g., Sankararaman et al., 2014; Vernot et al., 2016; Wall et al., 2013). Furthermore, some non-African populations have interbred with Denisovans, another archaic human, with levels of Denisovan introgression ranging from <1% in East and South Asians up to 5% in Oceanians and Southeast Asians (e.g., Browning, Browning, Zhou, Tucci, & Akey, 2018; Qin & Stoneking, 2015; Reich et al., 2010; Reich et al., 2011; Sankararaman, Mallick, Patterson, & Reich, 2016). Additionally, Browning et al. (2018) found that introgressed Neanderthal haplotypes in contemporary Europeans and East Asian genomes are equally similar to the Altai and Vindija Neanderthal genome sequences; thus, these authors argue that if there were multiple pulses of Neanderthal introgression, these pulses must originate from closely related Neanderthal populations. Two main models to explain regional differences in Neanderthal introgression have been proposed, which we will refer to as the “multiple pulses” and “dilution” models, respectively. The “multiple pulses” model was described by Vernot et al. (2016) based on whole genome sequences from the 1000 Genomes Project along with novel genome Oceanian genome sequences (it should be noted that this dataset does not include any Near Eastern populations). Vernot et al. (2016) found that Oceanians had less introgression than all other non-Africans in their dataset so they argued that Europeans, South Asians, and East Asians shared a second pulse of introgression to the exclusion of Oceanians. These authors also argued that East Asians received a third pulse of introgression, which was not found in the other studied regions.

An alternative, “dilution” model was put forth by Lazaridis et al. (2016) based on analyses of genomic sequence data from archaeological populations from the Levant and nearby regions along with SNP genotypes from contemporary, global populations. Lazaridis et al. (2016) and others (e.g., Rodriguez-Flores et al., 2016; Taskent et al., 2017) found that Near Eastern populations have less Neanderthal introgression than other Eurasians. Lazaridis et al. (2016) also found that Near Eastern populations have high levels of ancestry deriving from a group known as the “Basal Eurasians” that diverged from all other non-Africans prior to the separation of western Eurasians from eastern non-Africans (i.e., eastern Eurasians, Native Americans, and Oceanians). Basal Eurasian ancestry is highest in the Near East, lower in the rest of western Eurasia and South Asia, and absent elsewhere (Lazaridis et al., 2014; Lazaridis et al., 2016). Interestingly, Lazaridis et al. (2016) found that the amount of Basal Eurasian ancestry in ancient western Eurasians is strongly negatively correlated with Neanderthal introgression such that an individual with 100% Basal Eurasian ancestry would have little to no introgression, suggesting that Basal Eurasians did not receive any Neanderthal introgression. Thus, Lazaridis et al. (2016) posited that there was a single pulse from which virtually all Neanderthal introgression is derived. They argued that variations in levels of introgression could be largely attributed to dilution from populations with Basal Eurasian ancestry (Lazaridis et al., 2016). Not much else is known about the Basal Eurasian population, but it has been argued that they represent one or more populations indigenous to the Near East (Lazaridis et al., 2016; Rodriguez-Flores et al., 2016).

There is consensus that Neanderthal introgression first occurred in the Near East (Alves et al., 2012; Currat & Excoffier, 2011; Green et al., 2010; Lazaridis et al., 2016; Prüfer et al., 2014; Wall et al., 2013). Two particular regions of interest are the Levant and southern Arabia because they are hypothesized starting points for the OOA dispersal along a northern dispersal route (NDR) or southern dispersal route (SDR) route, respectively (e.g., Macaulay et al., 2005; Mirazón Lahr & Foley, 1994; Mirazón Lahr, 2016; Quintana-Murci et al., 1999; Rowold, Luis, Terreros, & Herrera, 2007). Presence of Neanderthals is well-documented in the Levant around 50-70 kya as Neanderthal fossils that date to 50- 70 kya and 48-60 kya have been recovered from sites such as Amud and Kebara, respectively, in what is now Israel (Valladas et al., 1987; Valladas et al., 1999). An early modern human calvaria was recovered from Manot Cave (also in what is now Israel) dating to around 55 kya, which suggests that modern humans and Neanderthals overlapped geographically and temporally in the Levant (Hershkovitz et al., 2015). There are also a number of archaeological sites in southern Arabia with Mode 3 (i.e., Middle Stone Age or Middle Paleolithic) stone artifacts ranging from around 120 to 55 kya, which may have been made by Neanderthals (Armitage et al., 2011; Delagnes et al., 2012; Delagnes, Crassard, Bertran, & Sitzia, 2013; Oppenheimer, 2012; Rose et al., 2011). The most recent of these sites is from Wadi Surdud in what is now Yemen and has been dated to around 55 kya (Delagnes et al., 2012). Given the archaeological evidence of Neanderthals in the Levant and southern Arabia and the genetic evidence that introgression occurred very soon after the main OOA dispersal, it can be assumed that introgression occurred within the Levant or southern Arabia (Fu et al., 2014; Sankararaman et al., 2012; Seguin-Orlando et al., 2014).

In this study, we compare levels of Neanderthal introgression in populations from the Levant and southern Arabia in order to investigate the first Neanderthal introgression event and to better understand human evolutionary history in the Near East. We conduct these analyses using genotyping data for >400,000 single nucleotide polymorphisms (SNPs) from two Neanderthal genomes, Levantine and southern Arabian populations, and comparative populations from around the world. In the first stage of the project, we calculate f_4_ and D statistics to estimate proportions of Neanderthal introgression and compare levels of introgression in Levantine and southern Arabian populations as well as in other populations from around the world. In the second stage of the project, we identify 1,581 putative Neanderthal-introgressed SNPs in our dataset. From these 1,581 SNPs, we compare Neanderthal allele frequencies in populations throughout the world to test if introgression is distributed similarly or differently throughout the genome. Based on levels of Neanderthal introgression and the distribution of introgression throughout the genome, we infer that introgression occurred prior to separation of Levantine and southern Arabian populations.

## MATERIALS AND METHODS

### Datasets analyzed

We analyzed genotyping data from the Affymetrix Human Origins array (Patterson et al., 2012) from three Yemeni populations (n=90) and three Eritrean individuals generated by Vyas, Al-Meeri, & Mulligan (2017a) along with populations from around the world selected from the “fully public” and “restrictive” datasets from Lazaridis et al. (2014) to yield a total modern human dataset of 2058 individuals from 165 populations. The data from Vyas et al. (2017a) are available on the Dryad Digital Repository (https://doi.org/10.5061/dryad.1pm3r) (Vyas, Al-Meeri, & Mulligan, 2017b). Data from the Altai Neanderthal, composite Vindija Neanderthal, and Denisovan genomes were also obtained from the “fully public” dataset from Lazaridis et al. (2014) (Green et al., 2010; Meyer et al., 2012; Prüfer et al., 2014; Reich et al., 2010). Data from Lazaridis et al. (2014) were converted to Plink format using EIGENSOFT and then merged with the data from Vyas et al. (2017a) using Plink v1.07 (Patterson, Price, & Reich, 2006; Price et al., 2006; Purcell et al., 2007). To avoid the effects of sex-biased demographic events, we only analyzed data from autosomal SNPs. This dataset had overlapping data for 462,712 SNPs and was used for f_4_ ratio calculations.

For D statistic calculation and identification of putative Neanderthal-introgressed SNPs, we included data for the inferred Human-Chimp ancestral alleles. These ancestral alleles were extracted from the 1000 Genomes Project VCF files using bcftools; output was filtered and then converted to Plink format using Python scripts (Li, 2011; The 1000 Genomes Project Consortium, 2015). High confidence ancestral allele data were recovered for 419,102 of the 462,712 available SNPs. These 419,102 SNPs were used for D statistic calculations. Putative Neanderthal-introgressed SNPs were also filtered and identified using this set of SNPs.

Throughout this study, the BedouinA, BedouinB, Druze, Jordanian, Lebanese, Palestinian, and Syrian populations from Lazaridis et al. (2014) were considered as representative Levantine populations, and the Yemen Desert, Yemen Highlands, and Yemen Northwest from Vyas et al. (2017a) as well as the Saudi and Yemenite Jewish populations from Lazaridis et al. (2014) were considered as representative southern Arabian populations; a map of these populations is depicted in Figure 1. Consistent with Vyas et al. (2017a), we labeled the Yemeni population from Lazaridis et al. (2014) as “Kuwaiti Yemeni” and did not include them as a southern Arabian population.

**Figure 1.**
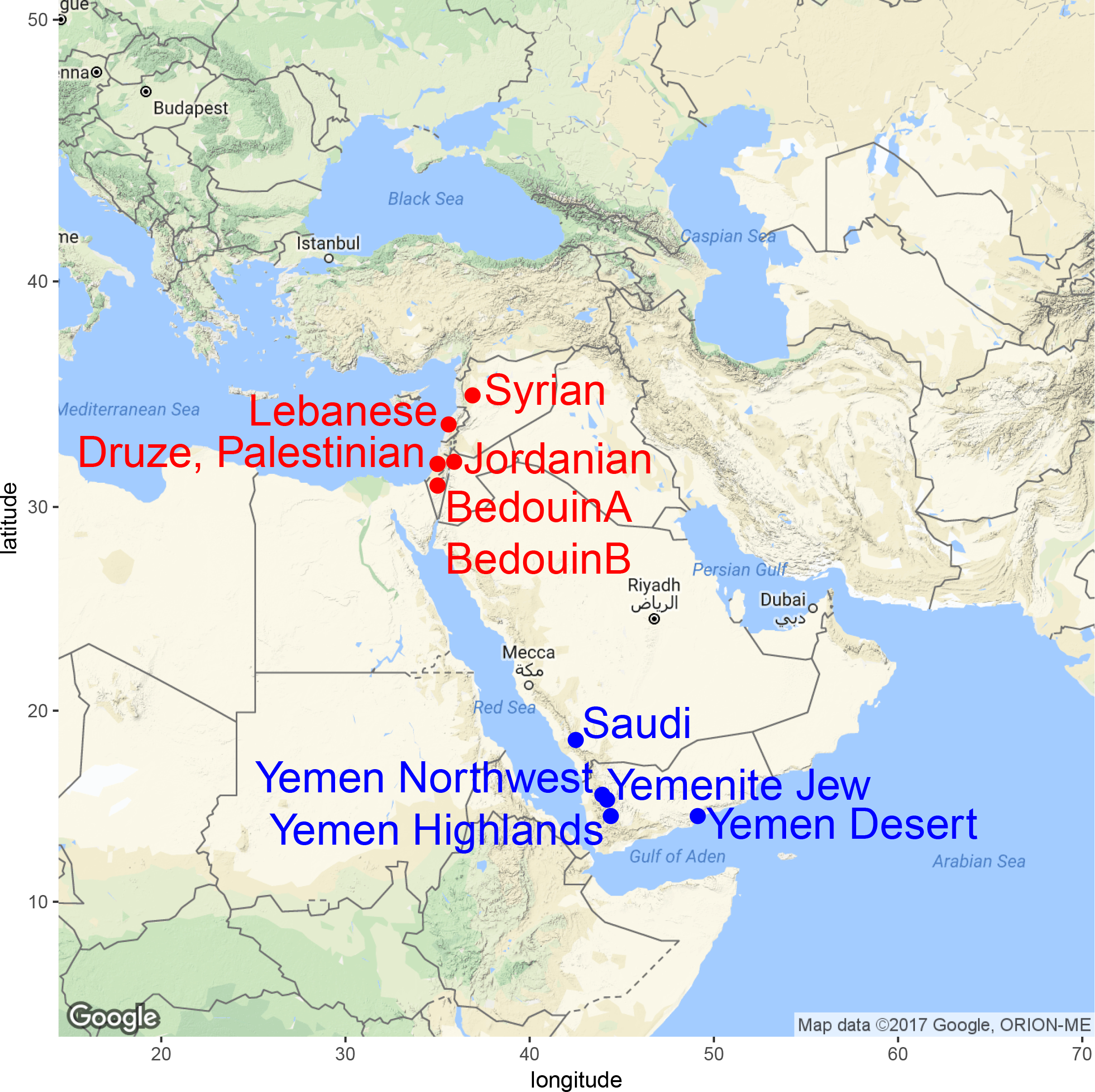
Map showing Levantine and southern Arabian populations. Levantine populations are labeled in red, and southern Arabian populations are labeled in blue. The latitude and longitude coordinates were provided by Lazaridis et al. (2014) and GeoNames (http://www.geonames.org/).

All populations analyzed in this study are listed in Table S1 along with their sample sizes, geographic region identifiers, and the analyses in which they were used. Populations were grouped into regions based on geography (e.g., Oceania, Siberia, West Africa), and some regions were also grouped into larger regions (e.g., eastern Europe, western Europe, northern Europe, and southern Europe were grouped into Europe); groupings were assigned based on groupings from Vyas et al. (2017a), which were derived from Lazaridis et al. (2014). In many analyses, populations were also pooled based on regions or larger regions (e.g., Europe, South Asia, Oceania) in order to provider clearer comparisons between different parts of the world; these pooled populations are listed in Table S1.

### Calculation of f_4_ and D statistics

We used the *qpF4Ratio* and *qpDstat* programs in the software package Admixtools to calculate f_4_ ratios and D statistics, respectively, to estimate Neanderthal introgression (Patterson et al., 2012). In order to estimate proportions of Neanderthal introgression, we calculated f_4_ ratios of the form of *f_4_(Altai Neanderthal, Denisovan; X, Mbuti)/ f_4_(Altai Neanderthal, Denisovan; Vindija Neanderthal, Mbuti)*, which estimates the excess frequency at which X (i.e. a test population) shares alleles with the Altai Neanderthal relative to the frequency at which the non-admixed Mbuti share alleles with the Altai Neanderthal (Patterson et al., 2012; Prüfer et al., 2014). The value of this ratio is the estimated proportion of introgression into population X. In addition to testing individual populations, ratios were also calculated for the regionally-defined pooled populations due to the fact that there is variability in levels of Neanderthal introgression within geographic regions (e.g., not all European populations have the same level of introgression) (Vernot et al., 2016). By comparing proportions of Neanderthal introgression in the Levantine and southern Arabian populations to the pooled populations, broader comparisons that summarize regional variations could be made. The f_4_ ratio values for the Levantine and southern Arabia populations were then compared to each other and to values for other individual and pooled populations in the dataset.

Due to the large standard errors associated with f_4_ ratios, D statistics were also used to compare the levels of Neanderthal introgression between pairs of populations (Green et al., 2010; Prüfer et al., 2014). Unlike f_4_ ratios, D statistics provide a direct comparison of the level of introgression between populations X and Y to test if there is a difference; mathematically, D statistics compare the number of derived SNPs shared between the Altai Neanderthal and population X to the number of derived alleles shared between the Altai Neanderthal and population Y (Durand, Patterson, Reich, & Slatkin, 2011; Green et al., 2010; Patterson et al., 2012). We calculated D statistics of the form *D(Ancestral, Altai Neanderthal; Y, X)* where X and Y represent different Levantine/southern Arabian populations for all 66 combinations of the seven Levantine and five southern Arabian populations in order to compare levels of Neanderthal introgression between individual Levantine and southern Arabian populations; for example, *D(Ancestral, Altai Neanderthal; Druze, Palestinian)* compared levels of Neanderthal introgression between the Druze and Palestinians. D statistics of this form were also calculated to compare the levels of introgression in the Levantine and southern Arabian pooled populations (Y) to other pooled populations (X) in the dataset to test for differences in levels of introgression [e.g., *D(Ancestral, Altai Neanderthal; Levant, Europe)* compared levels of Neanderthal introgression between Levantine and European populations.

The standard error of a D statistic follows an approximately normal distribution (Durand et al., 2011). Thus for this study, we used a cut-off of |Z| > 2 to determine whether a D statistic was significant or non-significant as this corresponded to an approximately 95% confidence interval. If Z < −2, the D statistic was significantly negative and Population X had less introgression than Population Y. Alternatively, if Z > 2, the D statistic was significantly positive and Population X had more introgression than Population Y.

Oceanian populations (i.e., Australian, Bougainville, and Papuan) were excluded from all f_4_ and D statistic calculations due to their significant levels of Denisovan introgression (Reich et al., 2011). The f_4_ ratio used above [*f_4_(Altai Neanderthal, Denisovan; Oceanian, Mbuti)/ f_4_(Altai Neanderthal, Denisovan; Vindija Neanderthal, Mbuti)*] would be confounded by the significant levels of Denisovan introgression found in these populations and would make it appear as if the populations had minimal or no archaic introgression (Prüfer et al., 2014). Additionally, D statistics would exaggerate the amount of Neanderthal introgression found in these populations due to the similarity of Neanderthal and Denisovan introgression (Prüfer et al., 2017). To confirm that other populations did not have detectable levels of Denisovan introgression, we calculated D statistics of the form of *D(Ancestral, Denisovan; Mbuti, X)* where X represents each of the populations in the dataset (Meyer et al., 2012; Reich et al., 2010). These statistics tested for the presence of Denisovan introgression; values that were significantly positive (Z > 2) would indicate that detectable levels of Denisovan introgression were present.

### Identification of Neanderthal-introgressed SNPs

All previously published analyses that compare the distribution of Neanderthal introgression in the genome were performed using data from whole genome sequences and inferred introgressed Neanderthal haplotypes (e.g., Sankararaman et al., 2014; Sankararaman et al., 2016; Vernot & Akey, 2014; Vernot et al., 2016). We created a novel method to compare how Neanderthal introgression is distributed throughout Levant and southern Arabian genomes using the existing SNP genotyping data from the Affymetrix Human Origins array. We designed a pipeline to identify introgressed SNPs that had introgressed from Neanderthals into modern humans based on the SNPs in the Affymetrix Human Origins dataset. Using maps of putative Neanderthal haplotypes from Vernot & Akey (2014) or Vernot et al. (2016), we filtered the 419,102 SNPs (where we have genotyping data from the Neanderthal genomes and the inferred Human-Chimp ancestral alleles) down to only SNPs occurring on the introgressed haplotypes (Table S2). Filtration was run in parallel based on the haplotype maps from Vernot & Akey (2014) and Vernot et al. (2016). In order to filter out SNPs that were polymorphic within Neanderthals (and thus remove any ambiguity of which allele was representative of Neanderthals), we only included sites where both the Altai Neanderthal and Vindija Neanderthal genomes were homozygous for the same allele. Next, in order to only retain SNPs where Neanderthals had the derived allele, SNPs where the Neanderthal allele matched the inferred Human-Chimp ancestral allele were further filtered out. A total of 18,081 SNPs from the Vernot & Akey (2014) map and 11,192 SNPs from the Vernot et al. (2016) map remained.

Finally, to confirm that a SNP was due to Neanderthal introgression and not due to an ancient polymorphism shared between Neanderthals and modern humans, SNPs were removed if the Neanderthal alleles were present in sub-Saharan African populations (since Neanderthal introgression is not thought to have occurred in Africa). Since many sub-Saharan African populations have low levels of Neanderthal introgression due to admixture from back-migrations and gene flow with Eurasia (e.g., Hodgson, Mulligan, Al-Meeri, & Raaum, 2014; Llorente et al., 2015; Pagani et al., 2012; Pagani et al., 2015; Skoglund et al., 2017; S. Wang, Lachance, Tishkoff, Hey, & Xing, 2013), we only filtered using African populations that were non-admixed per the ADMIXTURE results from Vyas et al. (2017a) (i.e., BantuKenya, Biaka, Dinka, Esan, Hadza, Luhya, Luo, Mende, Mbuti, and Yoruba). These ten sub-Saharan African populations have a total of 167 genotyped individuals and all of the SNPs with a non-zero frequency in this set of individuals were removed (i.e., if even one of these individuals had an allele matching the Neanderthal, the SNP was removed). Ultimately, we retained 1,388 SNPs from the Vernot & Akey (2014) map and 627 SNPs from the Vernot et al. (2016) map, which totaled 1,581 putative Neanderthal-introgressed SNPs; the numbers of SNPs retained at each level of filtration are presented in Table S2. The program ANNOVAR was used to identify the types of genomic regions in which the introgressed SNPs occurred (e.g., introns, exons, intergenic regions, UTRs) (K. Wang, Li, & Hakonarson, 2010). ANNOVAR was run on these 1,581 SNPs as well as the 419,102 SNPs from which they were filtered (to provide a comparison).

We calculated the frequencies of Neanderthal alleles in eight of the pooled populations (specifically, the Levant, southern Arabia, the Caucasus, East Asia, Europe, Siberia, Oceania, and South Asia). Furthermore, we conducted an association test to test if Neanderthal allele frequencies significantly differed between the Levant and southern Arabia.

Introgressed SNP filtration, allele frequency calculations, and the frequency association test described above were all completed using Plink v1.90b3.39 (https://www.cog-genomics.org/plink2) (Purcell et al., 2007). Additionally, all figures in this paper were generated using R v3.3.2 (R Core Team, 2016) and a variety of packages including ggplot2 (Wickham, 2009), ggmap (Kahle & Wickham, 2013), and cowplot (https://github.com/wilkelab/cowplot).

## RESULTS

### Neanderthal introgression estimates using f_4_ and D statistics

We estimated proportions of Neanderthal introgression using f_4_ ratios of the form of *f_4_(Altai Neanderthal, Denisovan; X, Mbuti)/ f_4_(Altai Neanderthal, Denisovan; Vindija Neanderthal, Mbuti)*, where X represents the test populations. Proportions of introgression for all seven Levantine, five southern Arabian, and twelve pooled populations from other regions are presented in Figure 2 and Table S3 and introgression proportion estimates for all 161 analyzed populations are presented in Table S4. The Levantine and southern Arabian populations cluster closely together indicating that they have similar levels of Neanderthal introgression as depicted in Figure 2.

**Figure 2.**
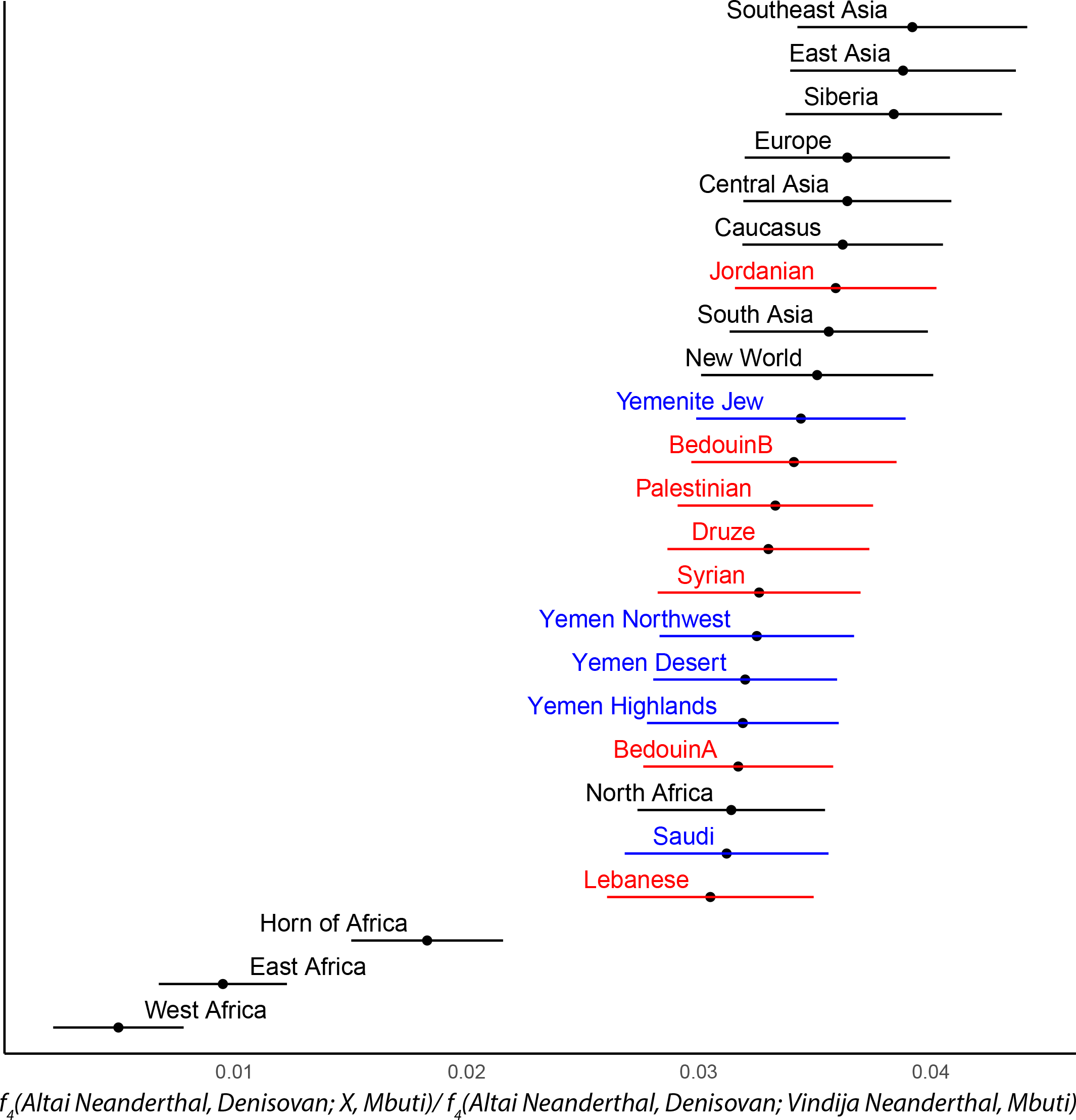
Estimated proportions of Neanderthal introgression based on f_4_ ratios. Plot of f_4_ ratios of Neanderthal introgression for all individual Levantine and southern Arabian populations and pooled populations from other regions. The f_4_ ratios estimate the proportion of Neanderthal introgression found in each tested population. Levantine populations are colored in red, southern Arabians in blue, and pooled populations in black. Error bars represent ±1 SE; f_4_ ratios are reported in Table S3.

Since f_4_ ratios have very large standard errors (i.e., see the large bars in Figure 2), we used D statistics as an additional measure to compare levels of introgression in different populations and to test if the differences in introgression between populations were statistically significant. We first calculated D statistics to compare the levels of Neanderthal introgression between individual Levantine and southern Arabian populations. Values for all 66 combinations are presented in Table S5 and values for the 35 combinations that compare a Levantine to a southern Arabian population are depicted in Figure 3. From the 66 population combinations, 63 were found to have non-significant values (−2 < Z < 2), which indicates that both regions have similar levels of introgression. From the three significant values, we find that Yemenite Jews have more Neanderthal introgression than the Highland Yemeni and the BedouinA and that Palestinians have more introgression than the BedouinA. We also calculated D statistics of the form *D(Ancestral, Altai Neanderthal; Y, X)* to compare levels of Neanderthal introgression in Levantine and southern Arabian populations (Y) to other populations (X) (Figure 4, Table S6). In Figure 4, it is notable that very few of the D statistics have error bars that overlap with 0, which indicates that most differences are statistically significant. We find that the Levantine and southern Arabian pooled populations have significantly more introgression than all sub-Saharan African pooled populations (i.e., East Africa, West Africa, and Horn of Africa) as the D statistics are significantly negative (Z < −2). Furthermore, the Levantine and southern Arabian pooled populations have significantly less introgression than most other non-African pooled populations as most of the D statistics are significantly positive (Z > 2). Consistent with the lack of differences in Neanderthal introgression between individual Levantine and southern Arabian populations that we reported above (Figure 3), we also find that the Levantine and southern Arabian pooled populations have very similar comparisons with other pooled populations (Figure 4).

**Figure 3.**
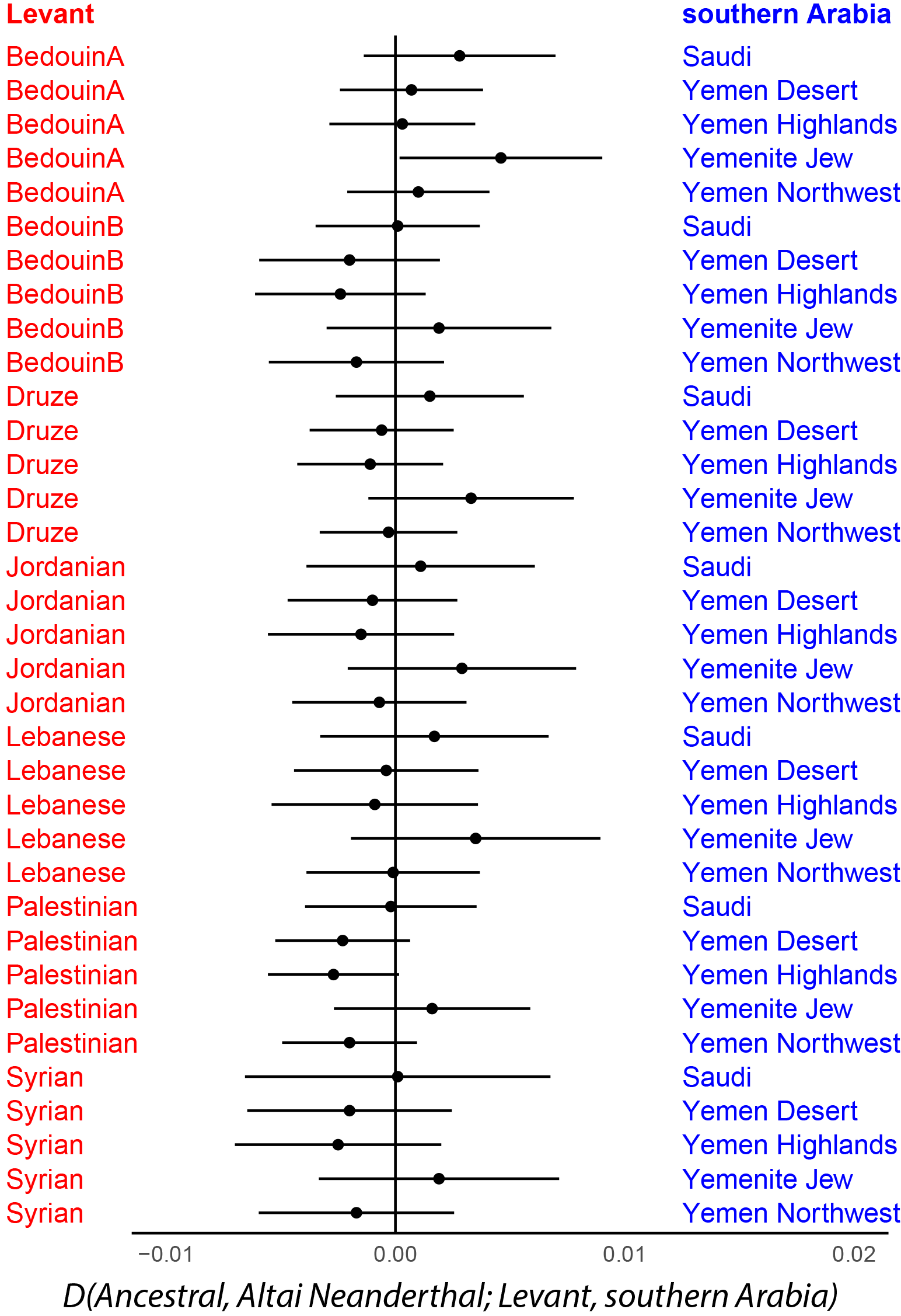
Comparisons of the level of Neanderthal introgression in individual Levantine and southern Arabian populations. Plot of D statistics for all 35 combinations of Levantine and southern Arabian populations. Positive D statistics indicate the southern Arabian population has more introgression, while negative D statistics indicate the Levantine population has more introgression. D statistics that are greater than two standard errors from 0 are considered statistically significant. Error bars represent ±2 SE; D statistics are reported in Table S5.

**Figure 4.**
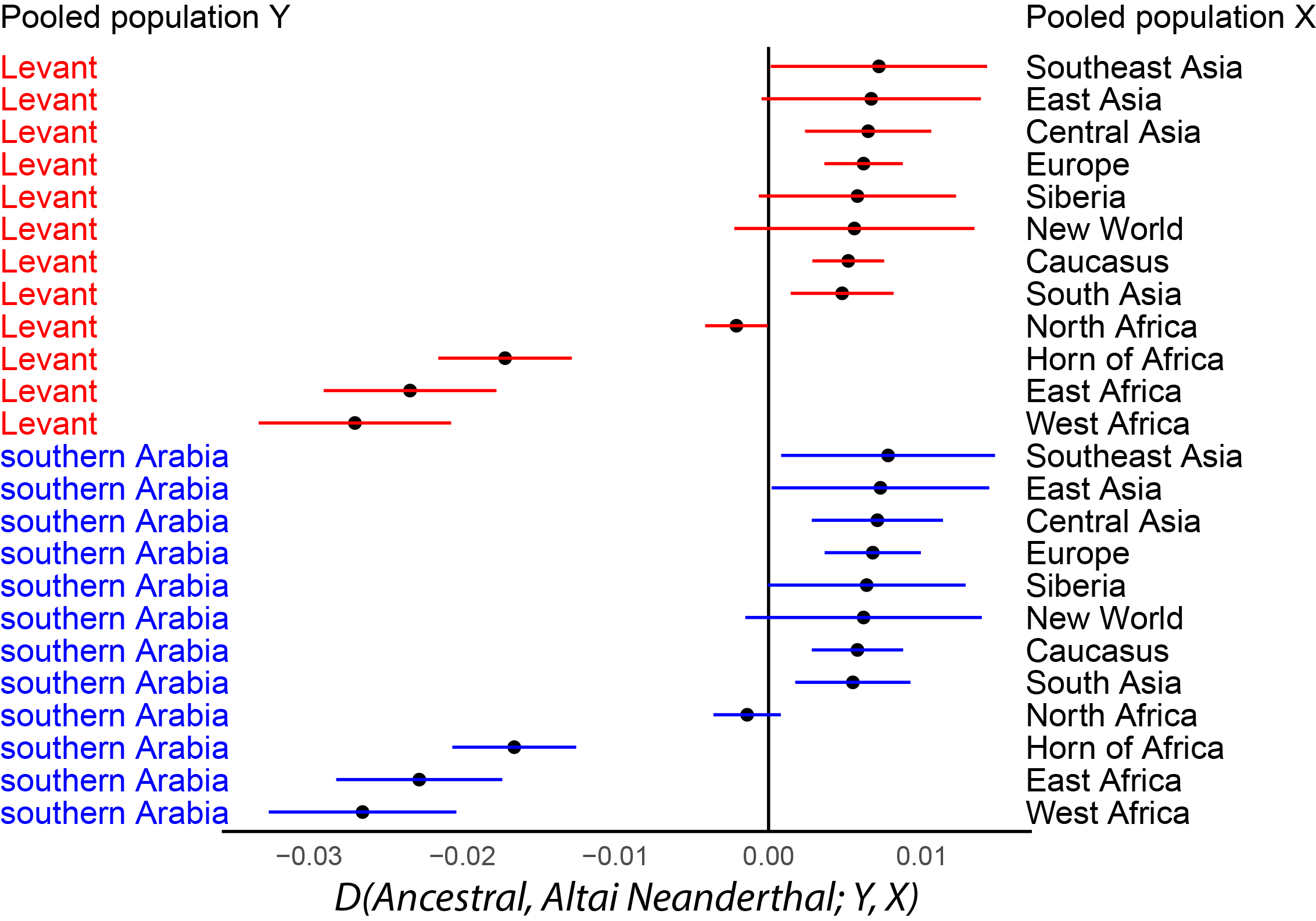
Comparisons of the level of Neanderthal introgression between Levantine and southern Arabia populations with other populations from around the world. Plot of D statistics of the form of *D(Ancestral, Altai Neanderthal; Y, X)* where X is one of twelve global pooled populations and Y is either the Levantine or southern Arabian pooled population. All comparisons with the Levant are colored in red and with southern Arabia in blue. Positive D statistics indicate X has more introgression, while negative D statistics indicate Y has more introgression. D statistics that are greater than two standard errors from 0 are considered statistically significant. Error bars represent ±2 SE; D statistics are reported in Table S6.

As the presence of Denisovan introgression can confound measures of Neanderthal introgression, all 164 populations were tested for detectable levels of Denisovan introgression (relative to the un-introgressed Mbuti) using D statistics of the form *D(Ancestral, Denisovan; Mbuti, X)* where X is the test population; these results are presented in Table S7. No populations outside of Oceania have detectable levels of Denisovan introgression (Z > 2), which is consistent with previous findings (Reich et al., 2011). Seven sub-Saharan African populations have significantly negative values (Z < −2), which indicate that they had more ancestral alleles (not present in the Denisovan) than derived alleles (shared with the Denisovans).

### Analysis of Neanderthal allele frequencies

From the dataset of 419,102 SNPs available for both Neanderthal genomes and the inferred human-chimp ancestral allele, we identified 1,581 putative Neanderthal-introgressed SNPs. These are SNPs where the derived allele is monomorphic in both the Altai and Vindija Neanderthal genomes and is not present in non-admixed Africans, which together strongly suggests that these SNPs are present in modern humans due to Neanderthal introgression. We analyzed these SNPs to understand how they are distributed throughout the genome using the program ANNOVAR (K. Wang et al., 2010). The introgressed SNPs are distributed across all 22 autosomes and across different types of functional regions (Table S8). The functional regions in which the introgressed SNPs occur generally reflect the 419,102 SNPs from which they were filtered (i.e., over 90% occur in intergenic and intronic regions; Table S8). Only 28 of the SNPs are exonic, and only 16 of these are nonsynonymous. Based on the SIFT scores (which identify if amino acid substitutions are tolerable or deleterious), the Neanderthal allele is deleterious for only one of the nonsynonymous SNPs (rs142330429 in the MGP gene) (Ng & Henikoff, 2003; K. Wang et al., 2010).

We calculated allele frequencies for the putative Neanderthal-introgressed SNPs in eight pooled populations (the Levant, southern Arabia, the Caucasus, East Asia, Europe, Siberia, Oceania, and South Asia). It is important to note that not all of the Neanderthal alleles are present in every part of the world, i.e. for any given introgressed SNP, the ancestral allele may be fixed in some regions but not in others. The number of Neanderthal alleles present in each pooled population at different frequencies is presented in Table S9. Comparisons of Neanderthal allele frequencies in different parts of the world are depicted in Figures 5 through 7. Figure 5 shows Neanderthal allele frequencies in southern Arabia versus in the Levant and Figures 6 and 7 depict Neanderthal allele frequencies in the Levant and southern Arabia, respectively, versus those in the Caucasus, Europe, South Asia, East Asia, Siberia and Oceania. The Neanderthal allele frequencies are very similar between the Levant and southern Arabia as the majority of points plot very close the Y=X line, indicating identical frequencies in both regions (Figure 5). Frequencies are still fairly similar when comparing the Levant or southern Arabia to the Caucasus, Europe, and South Asia, as points still plot close to the Y=X line (Figures 6 a-c and 7 a-c). However, very few points plot along the Y=X line when comparing the Levant or southern Arabia to East Asia, Siberia, and Oceania suggesting different Neanderthal alleles frequencies in these regions (Figures 6 d-f and7 d-f). Similarity of Neanderthal allele frequencies between the Levant and southern Arabia is also illustrated by the similarity of Figures 6 and 7.

**Figure 5.**
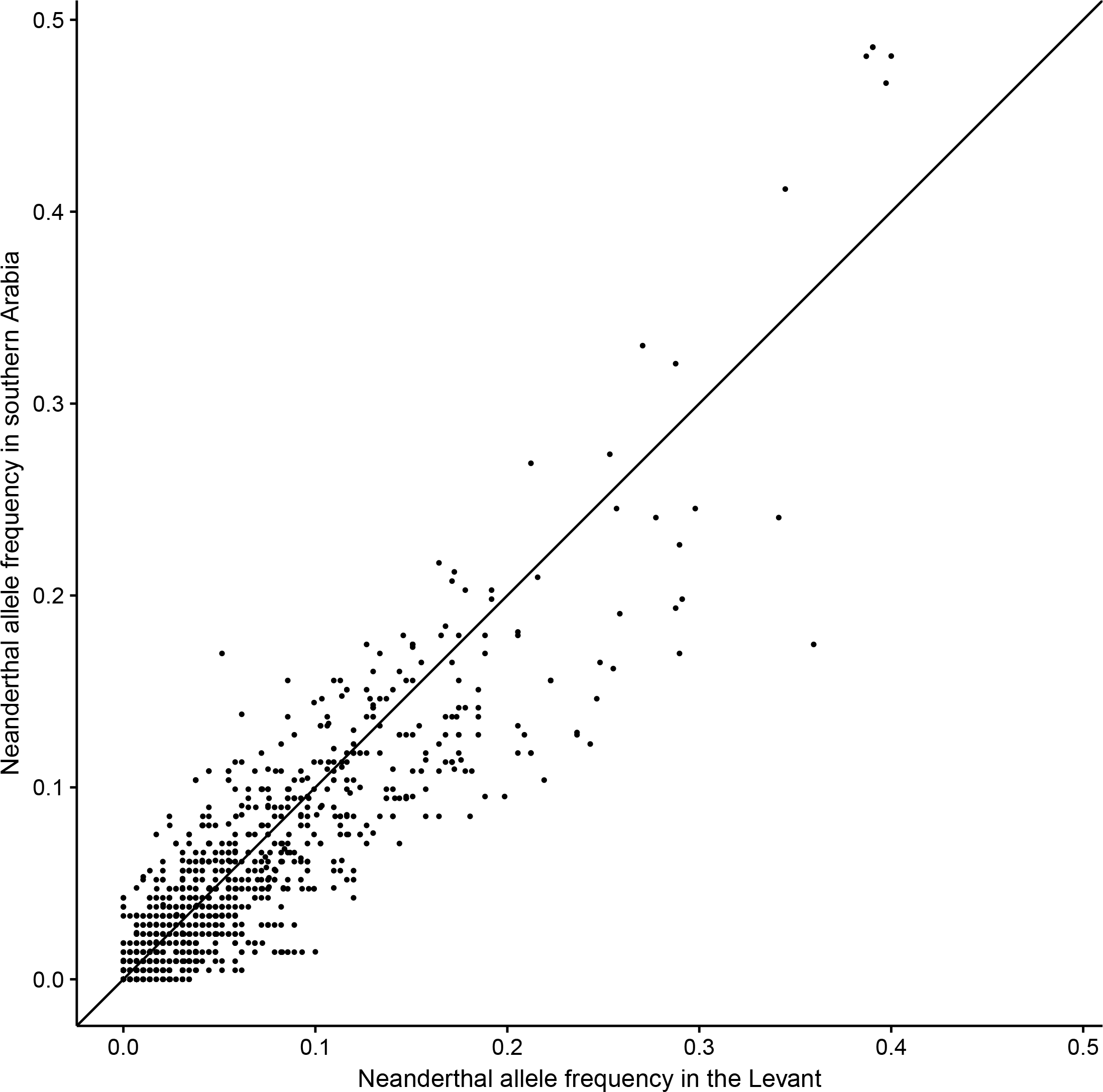
Comparisons of Neanderthal allele frequencies in the Levant and southern Arabia. Plot of Neanderthal allele frequencies in the Levant and southern Arabia. Each point represents one of the SNPs with the Neanderthal allele frequency in the Levant on the X-axis and its frequency in southern Arabia on the Y-axis. A point near the Y=X line indicates that the represented SNP has similar allele frequencies in the Levant and southern Arabia.

**Figure 6.**
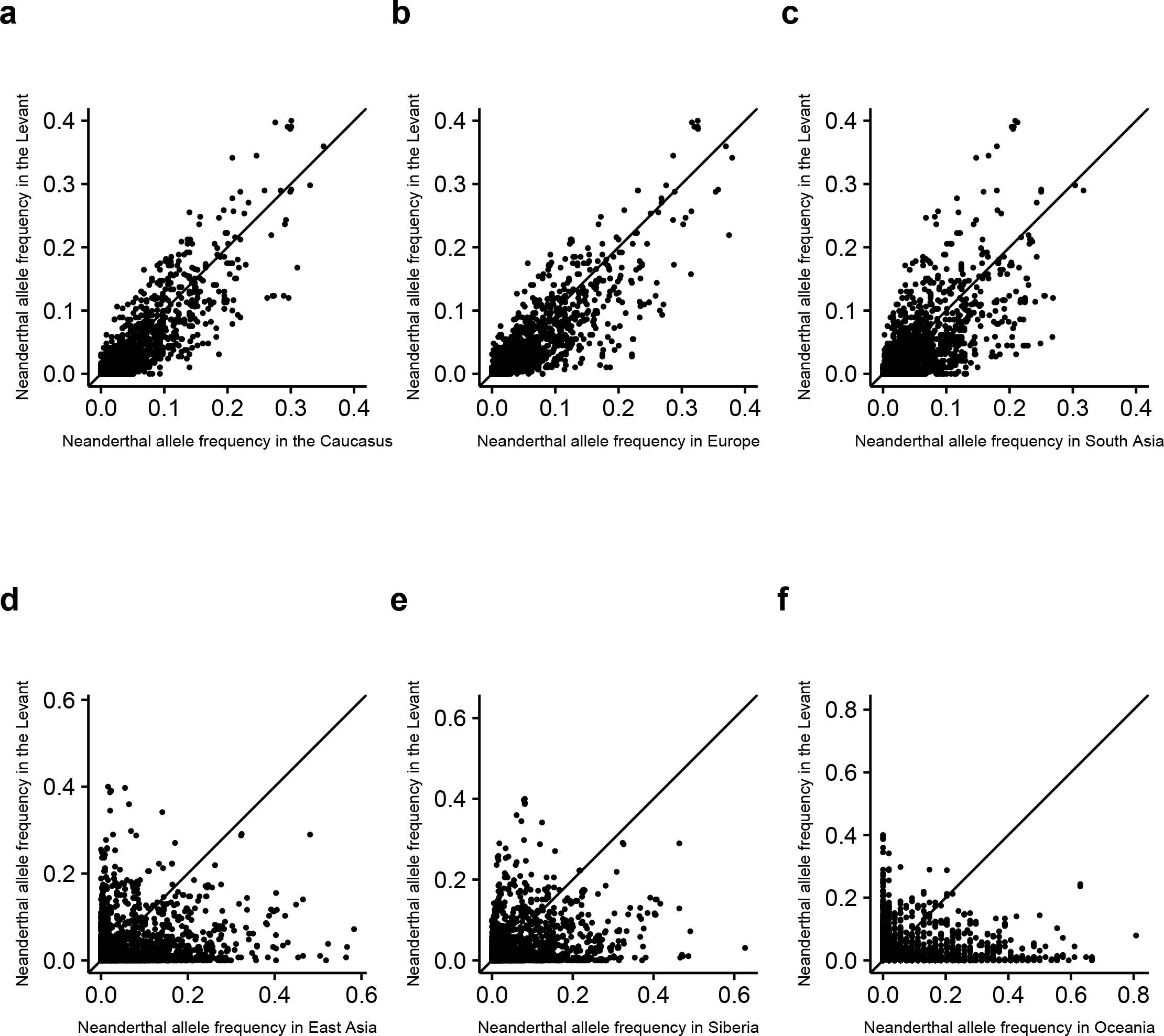
Comparisons of Neanderthal allele frequencies in the Levant and other regions of the world. Plot of Neanderthal allele frequencies in the Levant and six comparative regions of the world. Each point represents one of the SNPs with the Neanderthal allele frequency in the comparative region on the X-axis and the frequency in the Levant on the Y-axis. A point near the Y=X line indicates that the represented SNP has similar frequencies in the Levant and the other region. a) the Caucasus, b) Europe, c) South Asia, d) East Asia, e) Siberia, and f) Oceania. Note that the scale of axes differ across plots.

**Figure 7.**
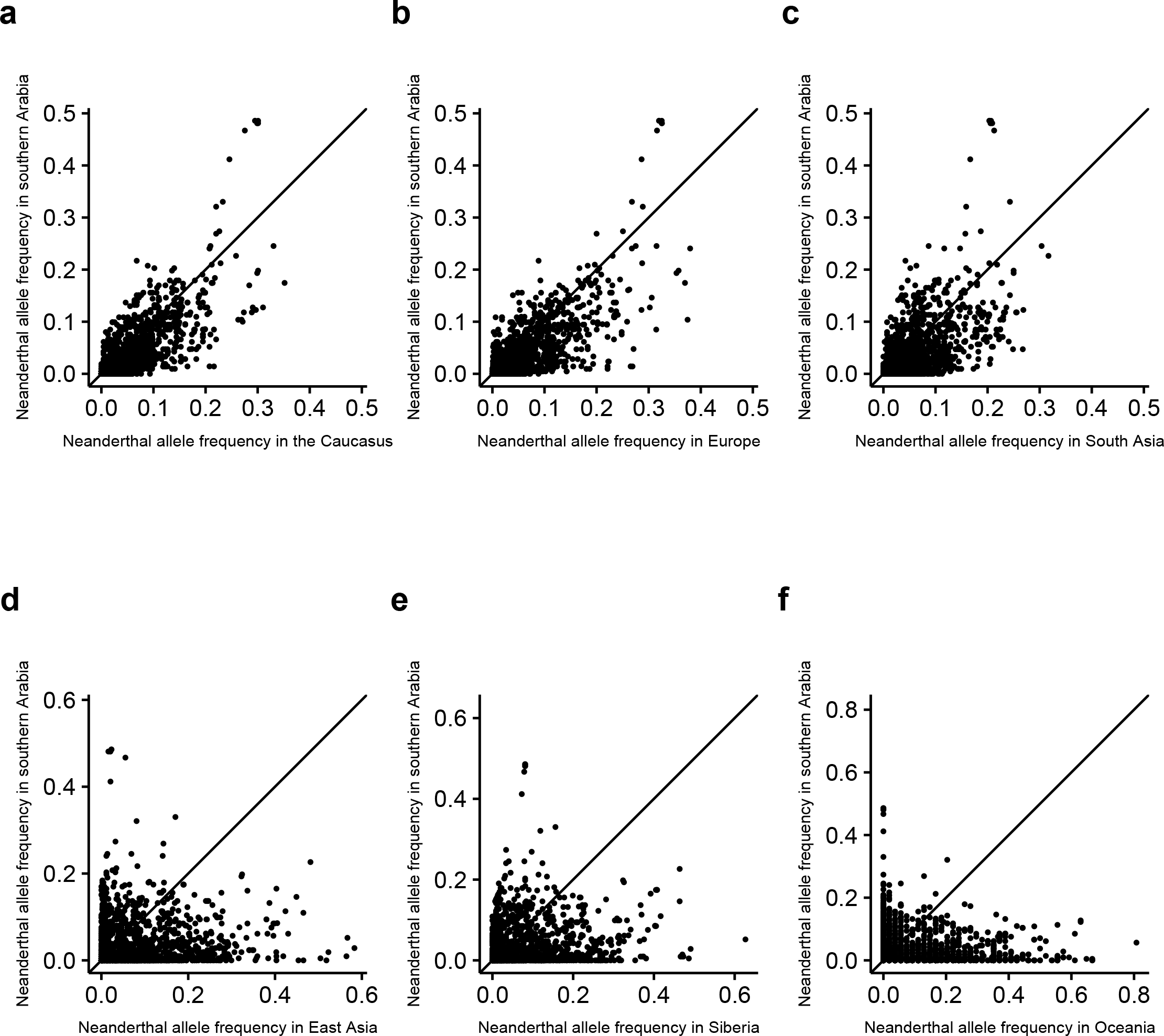
Comparisons of Neanderthal allele frequencies in southern Arabia and other regions of the world. Plot of Neanderthal allele frequencies in the southern Arabia and six comparative regions of the world. Each point represents one of the SNPs with the Neanderthal allele frequency in the comparative region on the X-axis and the frequency in southern Arabia on the Y-axis. A point near the Y=X line indicates that the represented SNP has similar frequencies in southern Arabia and the other region. a) the Caucasus, b) Europe, c) South Asia, d) East Asia, e) Siberia, and f) Oceania. Note that the scale of axes differ across plots.

We performed an association test to test whether any of the Neanderthal allele frequencies were significantly different between the Levant and southern Arabia. The p-values for each SNP along with allele frequencies in the Levant and southern Arabia and genomic information from ANNOVAR are presented in Table S10. Out of the 1581 SNPs, 309 are fixed for the ancestral allele (i.e., the Neanderthal allele is absent) in both regions meaning there are no testable differences for these SNPs. Of the remaining 1272 SNPs, only two SNPs had significant frequency differences after Bonferroni correction (p < 0.05/1272 = 3.93 × 10^−5^). These two SNPs (rs13091098 and rs4128294) are intronic and intergenic, respectively, meaning they are non-coding SNPs and the frequency differences are likely due to genetic drift, gene flow, or hitchhiking as opposed to direct selection. Overall, these results indicate that Neanderthal alleles have very similar frequencies in the tested Levantine and southern Arabian populations.

## DISCUSSION

In this study, we analyze Neanderthal introgression in the Levant and southern Arabia in order to investigate where introgression may have first occurred and how introgressed alleles may have dispersed throughout the Near East. Using >400,000 SNPs, we estimate the level of introgression in the Levant and southern Arabia and compare levels of introgression within these two regions and to other parts of the world. Also from this dataset of >400,000 SNPs, we identify and analyze 1,581 putative Neanderthal-introgressed SNPs to compare how introgression is distributed throughout the genome in different regions of the world.

### Comparing levels of Neanderthal introgression

We calculated f_4_ ratios to estimate proportions of Neanderthal introgression in the genome and D statistics to test if levels of introgression in two populations were statistically significantly different. Based on f_4_ ratios, we find that proportions of introgression are relatively low and very similar among the Levantine and southern Arabian populations (Figure 2; Table S3). These findings are consistent with previous findings that Near Eastern populations have lower levels of introgression than other non-Africans (Lazaridis et al., 2016; Rodriguez-Flores et al., 2016; Taskent et al., 2017). The low levels of introgression in these populations cannot be attributed to gene flow from sub-Saharan Africa as some of these Levantine and southern Arabian populations (such as the Druze and Saudis) were found to have minimal sub-Saharan ancestry (Vyas et al. 2017a).

While f_4_ ratios can provide estimates of Neanderthal introgression in individual populations, D statistics are used to test for statistically significant differences in Neanderthal introgression between pairs of populations. We calculated D statistics to compare the level of Neanderthal introgression between each of the Levantine and southern Arabian populations. These comparisons indicate that the Levantine and southern Arabian populations have statistically indistinguishable levels of introgression. From 66 combinations of Levantine/southern Arabian population pairs, 63 of the comparisons are non-significant (Figure 3; Table S5). Furthermore, the three significant differences have marginally significant Z-scores (ranging from 2.08 to 2.16), and only one demonstrates a difference between the Levant and southern Arabia (i.e., the Yemenite Jews have more introgression than the BedouinA). We also calculated D statistics to compare the level of Neanderthal introgression in the Levantine and southern Arabian pooled populations (Y) to other pooled populations (X) (Table S6). The Levantine and southern Arabians have significantly more introgression than sub-Saharan Africans as depicted by significantly negative D statistics in Figure 4. Furthermore, the Levant and southern Arabia have significantly less introgression than most other non-Africans as depicted by the significantly positive D statistics in Figure 4; these findings are consistent with previous studies of Near Eastern populations (Lazaridis et al., 2016; Rodriguez-Flores et al., 2016; Taskent et al., 2017). The Levantine and southern Arabian pooled populations also share very similar D statistic values when compared to other non-African pooled populations (compare top and bottom of Figure 4; Table S6). All of these results firmly indicate that the Levant and southern Arabia do not have detectable differences in levels of introgression, suggesting they may share the same introgression event. The question then arises of whether this introgression derives from similar or different parts of the Neanderthal genome.

### Comparing frequencies of Neanderthal alleles

It is known that different regions of the world have different Neanderthal haplotypes at different frequencies despite having strikingly similar levels of introgression. For example, Vernot et al. (2016) found that Europeans and South Asians have similar levels of Neanderthal introgression, but there are different Neanderthal haplotypes found across these regions (Sankararaman et al., 2016; Vernot et al., 2016). In order to test whether introgression is similarly or differently distributed throughout Levantine and southern Arabian genomes, we generated a protocol to identify Neanderthal-introgressed SNPs in the Affymetrix Human Origins dataset.

We ultimately identified a set of 1,581 putative Neanderthal-introgressed SNPs, which are distributed across all 22 autosomes. These SNPs are almost exclusively noncoding and occur primarily in intergenic regions (50.16%) and in introns (40.61%) with less than 2% of the SNPs occurring in exons (Table S8). When designing the Human Origins array (from which all of these data were generated), Patterson et al. (2012) chose SNPs without respect to functional elements and did not emphasize genes or regions around them, which resulted in an excess of noncoding SNPs. As a result, many of the genes associated with well-understood cases of adaptive introgression (e.g., Ding et al., 2014; Huerta-Sanchez et al., 2014; Mendez, Watkins, & Hammer, 2012; Vernot & Akey, 2014) are not present in this dataset. While a small subset of the 1,581 introgressed SNPs that we identified are associated with Neanderthal selective sweeps found by Sankararaman et al. (2014), the SNPs are often in the flanking regions instead of the gene body (e.g., *BNC2*) (Table S10). One notable exception is *AKAP13* (i.e., a gene involved in the regulation of toll-like receptors, which regulate innate immunity) where there are multiple introgressed SNPs within both exons and introns, and the Neanderthal alleles are all in relatively high frequencies in both the Levant and southern Arabia (38-48%) (Table S10) (Sankararaman et al., 2014; Shibolet et al., 2007). The presence of multiple introgressed SNPs at high frequencies suggest that the Neanderthal alleles around *AKAP13* have been selected in the Levant and southern Arabia, consistent with previous findings of enrichment of archaic haplotypes around toll-like receptor genes (Dannemann, Andrés, & Kelso, 2016; Sankararaman et al., 2014).

With the exception of *AKAP13*, we assume the vast majority of the 1581 introgressed SNPs have been primarily influenced by genetic drift and gene flow and, therefore, can be used to reconstruct evolutionary history. Thus, for these 1,581 introgressed SNPs, we compared the Neanderthal allele frequencies in the Levant and southern Arabia to six other parts of the world to test for similarities or differences in allele frequencies. The Neanderthal alleles have very similar frequencies in the Levant and southern Arabia, which indicates that introgression is distributed similarly through the genome in these two geographic regions (Figure 5). This is further supported by association test results, which indicate that only two SNPs have significant differences in allele frequencies (Table S10). We also find that Neanderthal allele frequencies in other regions vary in relation to their geographic distances from the Levant/southern Arabia. In regions closer to the Near East, such as the Caucasus, Europe, and South Asia, Neanderthal frequencies remain relatively similar between both the Levant and southern Arabia and these regions (as points cluster along the Y=X line) (Figures 6a-c, 7a-c). However, in more distant regions (i.e., East Asia, Siberia, and Oceania), allele frequencies differ much more between Levantine or southern Arabian populations and these regions; for example, there are many SNPs where the Neanderthal allele is at high frequency in East Asia, Siberia, and Oceania yet is at low (or even 0%) frequency in the Levant and southern Arabian (Figures 6d-f, 7d-f). These results are consistent with an isolation-by-distance model, whereby gene flow with neighboring regions keeps frequencies similar and isolation from more distant regions allows frequencies to drift independently.

### Inferences about the history of introgression in the Levant and southern Arabia

Our findings suggest that Neanderthal introgression is strikingly similar and indistinguishable between populations in the Levant and southern Arabia, both in that the two regions have statistically indistinguishable levels of introgression and that they have equally similar frequencies of Neanderthal alleles. Furthermore, consistent with previous studies of Near Eastern populations, we find that the Levantine and southern Arabian populations have less introgression than most other non-Africans. In order to make inferences about where introgression first occurred or how it spread across the Near East, these overall findings need to be interpreted within the context of different models of introgression; specifically, we look at the “multiple pulses” model from Vernot et al. (2016) and the “dilution” model from Lazaridis et al. (2016).

Of the two models, it is much simpler to contextualize our results within the “multiple pulses” model. Within this model, Levantine and southern Arabian populations are proposed to have only one pulse of introgression. Scenarios can be constructed with early modern humans using either the NDR or SDR and then receiving a single pulse of introgression in the Levant or southern Arabia, respectively. The dispersing wave front would then continue to people the rest of the world (including the remainder of the Near East) and interbreed with Neanderthals again as Europe, South Asia, and East Asia were peopled. The additional pulses in Europe, South Asia, and East Asia would explain why Levantine and southern Arabian (and other Near Eastern) populations have less Neanderthal introgression than most other non-Africans. Furthermore, as Levantine and southern Arabian populations would both have only received one pulse, it would explain why they have the same levels of introgression. Under this model, our results are consistent with introgression first occurring either in the Levant along an NDR dispersal or in southern Arabia along an SDR dispersal. It is difficult to argue in favor of introgression occurring in one region over the other in part due to the overall similarity of introgression in the Levant and southern Arabia.

In terms of the “dilution” model, the means by which Levantine and southern Arabian populations have less introgression than other non-Africans yet maintain the same level of Neanderthal introgression and the same Neanderthal alleles as each other is more complicated. The dilution model posits that Neanderthal introgression will be diluted If an introgressed population (either from within the Near East or from elsewhere) admixed in the Near East with a Basal Eurasian population that carries no (or minimal) Neanderthal introgression. A scenario can be constructed wherein contemporary Levantine and southern Arabian populations derive from a common ancestral population, which peopled/re-peopled the Levant and southern Arabia (and other parts of the Near East). The ancestral population would have derived from a Basal Eurasian population that admixed with an introgressed population and thus would have some, but less, introgression than other non-Africans. This ancestral population would have spread from one region to the other (i.e., from southern Arabia to the Levant or vice versa). This population would have also replaced any non-introgressed populations in these regions, as significant amounts of admixture with non-introgressed populations would create differences in Neanderthal introgression between contemporary Levantine and southern Arabian populations (which are not present in our results). Under this scenario, it would not be clear where in the Near East introgression occurred due to the similar levels of introgression in contemporary Levantine and southern Arabian populations. An alternate scenario is that non-introgressed Basal Eurasian populations and other introgressed Eurasian populations admixed independently in both the Levant and southern Arabia (resulting in dilution of Neanderthal introgression in both regions). We cannot refute this hypothesis; however, it is less parsimonious, as it requires multiple admixture events between introgressed and non-introgressed populations that ultimately resulted in populations with similar levels of introgression and Neanderthal allele frequencies throughout the Levant and southern Arabia.

Due to the overall similarity of introgression in the Levant and southern Arabia, different models of introgression do not provide a clear answer to where introgression first occurred. Since levels of introgression and Neanderthal allele frequencies are so similar between the two regions, it is not possible to pin the site of initial introgression to either the Levant or southern Arabia. However, from these findings, we can make inferences about the history of Levantine and southern Arabian populations. We hypothesize that introgression is similar in the Levant and southern Arabia because contemporary Levantine and southern Arabian populations derive from a common ancestral population. This common ancestral population could represent an introgressed population from the initial introgression event that spread from the Levant to southern Arabian or vice-versa. Alternatively, this common ancestral population could represent a population with diluted introgression that spread across and peopled the Near East.

Consistent with previous findings, our results also indicate that significant gene flow must have occurred between the Levantine and southern Arabian populations (Al-Abri et al., 2012; Fernandes et al., 2012; Fernandes et al., 2015; Vyas et al., 2017a; Vyas et al., 2016). Regardless of the origin of introgression in the Levant and southern Arabia or the relatedness of the populations in this regions, it would not be expected that selectively-neutral SNPs would maintain such similar frequencies in both regions (Tables S8, S10). Gene flow between the Levantine and southern Arabian populations is necessary to explain the high similarity of Neanderthal allele frequencies. The overall similarity of Neanderthal introgression between the Levant and southern Arabia can be most adequately explained by contemporary Levantine and southern Arabia populations descending from the same ancestral population with ongoing gene flow between the two regions.

## Summary

In this study, we analyzed Neanderthal introgression in Levantine and southern Arabian populations using genotyping data from a collection of >400,000 SNPs typed in >2000 individuals. We used f_4_ and D statistics to compare levels of introgression in different geographic regions. We also identified putative Neanderthal-introgressed SNPs in order to compare distribution of introgression throughout the genome. From these analyses, we find that Levantine and southern Arabian populations have extremely similar levels of Neanderthal introgression, which are higher than sub-Saharan Africans and lower than other non-Africans. Additionally, we find that these populations have very similar frequencies of specific introgressed Neanderthal alleles. These results suggest that contemporary Levantine and southern Arabian populations derive from a common, introgressed ancestral population, as opposed to introgression flowing from one region to the other. Furthermore, the extreme similarity of Neanderthal allele frequencies in both regions provides evidence of significant gene flow between these two regions.

## ACKNOWLEDGEMENTS

Thanks are owed to the University of Florida Research Computing support staff for analytic assistance. This work was conducted with support of the National Science Foundation [BCS-1258965 to C.J.M and D.N.V.]. This work derives from the dissertation of D.N.V. The authors have no conflicts of interest.

